# An updated genotype classification system for Zika viruses

**DOI:** 10.1101/760066

**Authors:** Hanna Nebenzahl-Guimaraes, Pieter Libin, Victor Pimentel, Marta Pingarilho, Kristof Theys, Ana B. Abecasis

## Abstract

Despite the Zika virus’ rapid spread throughout the Americas in 2015, its phylogenetic classification is limited to either an ‘African’ or ‘Asian’ genotype. This classification no longer reflects the present genetic diversity of circulating strains and their geographic reach. Using 414 publicly available Zika virus genomes from 40 different countries, we created alignments of different genomic fragments with high phylogenetic signal and used these to construct multiple maximum likelihood trees (IQTree). We observed groups of strains from three major geographic regions that consistently cluster into monophyletic clades: African (AF), Asian (AS) and American (AM), the latter of which cluster into four further sub-clades: Caribbean (C), South American (SA1 and SA2) and Central American (CA). The inter-genetic distances of these clades are all significantly greater than their intra-genetic distances (p=0.05). A decision demonstrates that only five nucleotide positions are needed to correctly classify 90% of our dataset into the newly described genotypes.

---

The Zika virus caused a global public health emergency in 2015 after it traveled from French Polynesia to Brazil and subsequently spread rapidly throughout the Americas [1]. Despite it being reported in 86 countries and causing ~30,000 cases in Brazil alone, only two lineages have been defined in the literature: the “African” lineage, named after the continent where ZIKV emerged in 1947, and the “Asian” lineage, named after the continent in which it has been circulating during the latter part of the 20^th^ century [2].

This classification no longer reflects the present genetic diversity of circulating strains and ignores the virus’ recent evolutionary adaptations. These adaptations are however associated with serious neuropathologies and novel modes of transmission (i.e., mother to child, sexual and blood transfusion transmissions), and greatly contrast with the mostly asymptomatic or mildly symptomatic African and Asian strains [3]. Accurate genotypic correlates to such phenotypic adaptations matter in clinical decision-making, such as the prescription of appropriate treatments, and in tailoring vaccines to maximize their efficacies in different settings.

Harnessing the boom of ZIKV sequences published during these last few years, we propose a new genotype classification of ZIKV lineages that more accurately reflects their evolutionary dynamics and spatial circulation. To this end, we investigate the consistency of ZIKV sequence clusters in different genomic segments, which indicates the existence of new genotypes. We subsequently validated these clusters by comparing their intra and inter-cluster genetic diversity and using machine learning algorithms for cluster grouping. We finally identified a set of genomic positions that make up the signature for the proposed genotypes.

We collected all whole-genome sequences available from Genbank, resulting in a combined dataset of 414 ZIKV sequences. To allow for visual inspection of consistent clusters, we used Phylogeny Diversity Analyzer to create a subset of 109 genomes, which is representative of the genetic diversity of all Zika genomes in our dataset [4]. This down-sampled dataset includes sequences collected in 40 different countries between 2008 and 2018. We evaluated the phylogenetic signal in this dataset by investigating windows of different genomic sizes and different genetic regions, using the quartet puzzling algorithm, as implemented in TREE-PUZZLE [5]. We found that 4000bp fragments and the first three non-structural genetic regions (i.e., NS1, NS2 and NS3) exhibited the highest signal with an average of 84.4% (95% CI = 81.2 – 87.6). To check for consistency of clustering across genomic regions, we constructed codon-correct alignments of the most phylogenetically informative genomic fragments [6] and inferred maximum likelihood trees for each of the genomic regions [7]. The trees were rooted using the Spondweni virus (i.e., GenBank sequence DQ859064.1).

We observed groups of strains from three major geographic regions, i.e., Africa (AF), Asia (AS) and America (AM), that consistently clustered with support of >= 95% across all trees (Fig 1A; detailed trees with bootstrap support may be found in the Supplement) [8]. Furthermore, the AM cluster consistently grouped into four sub-clusters: Caribbean-US (C), South America (SA1 and SA2), and Central America (CA). Based on these clusters, we assigned 373 strains from our overall dataset of 414 to one of the three major monophyletic clades (AF, AS or AM). Sixteen of the unclassified strains prevailed from Oceania, whose cluster we considered too small to merit a sub-genotype of its own (Fig 1B). We further assigned 227 strains from the AM genotype (n=247) into one of the four monophyletic sub-genotypes (C, SA1, SA2 or CA) (Fig 1B). Their inter-group genetic distances, computed using the Tamura-Nei model, were all significantly greater (p=0.05, established using ANOVA) than their intra-group genetic distances (Table 1) [9].

**Fig 1.**
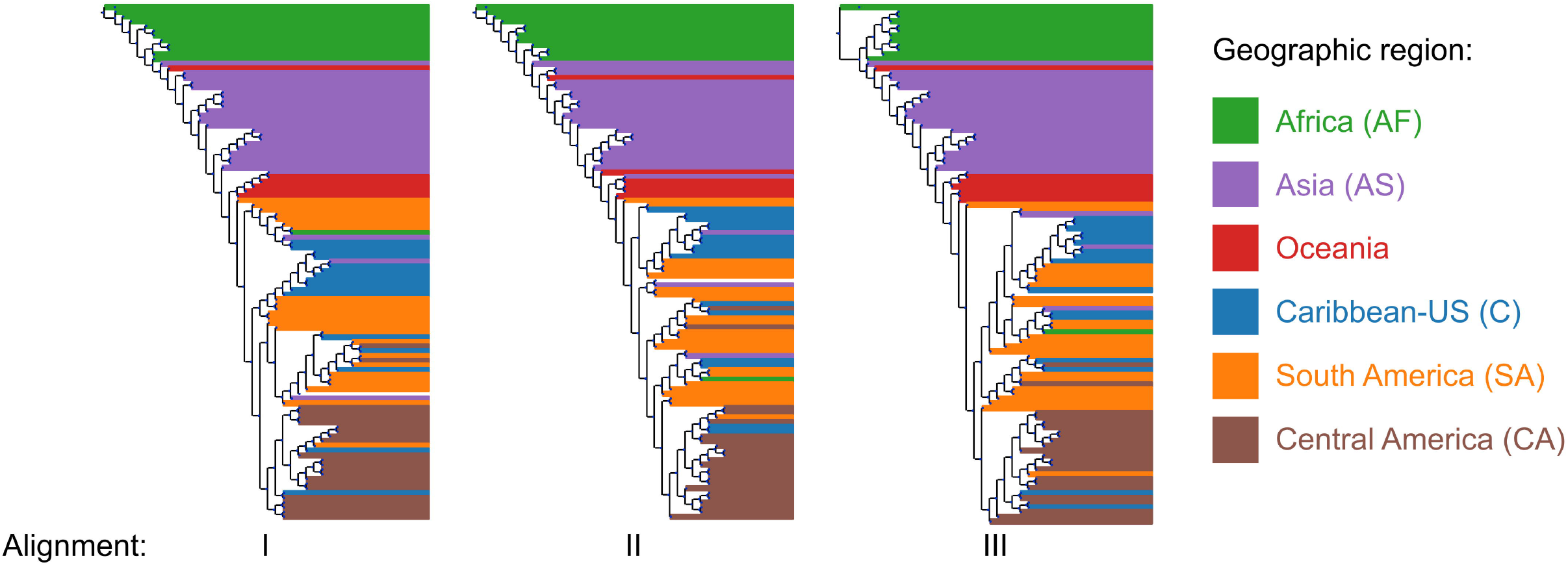
(A) Similarity and consistent clustering across maximum likelihood trees (IQTree) constructed using alignments: I (genome segment 1-4000bp), II (genome segment 4001-8000bp) and III (genes NS1-NS2). Color coding denotes geographic origin of strains. (B) Genotype-annotated maximum likelihood tree of whole genome sequences of full dataset (414 strains) using the GTR model in FastTree. Color-coding matches that of Panel A; highlighted areas of tree indicate proposed genotypes/sub-genotypes. Red stars indicate branch nodes that define major genotypes, yellow stars indicated branch nodes that define sub-genotypes (all of which have support > 0.85). (C) Decision tree output showing the five Amino Acid positions (numbered according to their coding within gene position in Reference sequence KJ77679) that successfully classify strains into Genotypes G1 - G3. The codon variant is highlighted in bold.

**Table 1.** Intra- and inter-genetic distances between proposed genotypes and sub-genotypes.

| Genotypes | AF | AS | AM |
| --- | --- | --- | --- |
| AF | **0.048** |  |  |
| AS | 0.125 | **0.008** |  |
| AM | 0.126 | 0.011 | **0.004** |

| Sub-genotypes | C | SA1 | SA2 | CA |
| --- | --- | --- | --- | --- |
| C | **0.002** |  |  |  |
| SA1 | 0.005 | **0.002** |  |  |
| SA2 | 0.005 | 0.004 | **0.003** |  |
| CA | 0.005 | 0.004 | 0.004 | **0.002** |

Legend: Numbers in bold denote intra-genetic distances.

A decision tree was estimated from phylogenetically informative sites (nucleotide variations at 5058 sites) using the C4.5 algorithm, 10-fold cross-validation and confidence factor of 0.25. This decision tree shows that the above-mentioned clusters can be explained using only five nucleotide positions (i.e., this tree correctly classifies 99.2% of the 373 strains). The positions that clearly stratify these clusters are a synonymous substitution in AA position 56 in gene C, a synonymous substitution in AA position 123 in NS1, a synonymous substitution in AA position 3 in NS2A, and nonsynonymous substitutions G99A in NS1 and M112V in NS5 (Fig 1C). The numbering of these positions is relative to reference sequence KJ77679 [10].

Based on our results, we propose the adoption of an expanded classification system for Zika viruses. Given that future outbreaks can change the spatial distribution of these genotypes, we propose not to name them according to geographic regions. Instead, we propose the usage of Genotype 1 (G1, formerly named African), Genotype 2 (G2, formerly named Asian) and Genotype 3 (G3, referred to above as AM). We further subdivide Genotype 3 in the following sub-lineages: 3a (G3a, referred to above as C), 3b (G3b, referred to above as SA1), 3c (G3c, referred to above as SA2), and 3d (G3d, referred to above as CA) (Fig 1C).

Due to the paraphyletic structure of the tree, some sequences could not be assigned to any genotype or sub-genotype. This is likely due to the dense sampling of sequences reflecting recent evolution of a virus that only recently became pandemic. As a result, we are capturing the complete genetic diversity of the viral strains before the establishment of clear genotype clusters, i.e. before actual ‘speciation’ into genotypes and sub-genotypes. This is opposite to what has been found in other viruses, where monophyletic clusters reflect a much more ancient evolutionary history and genotypes and sub-genotypes became defined a long time ago. In HIV, however, it has been shown that strains which were circulating in the earlier days of the epidemic are paraphyletic to established HIV-1 subtypes, which is in line with what we observe in our ZIKV phylogeny [11].

This updated classification system will serve as a reference and starting point for future classification expansions, when future outbreaks cause a further increase of the genetic diversity of the virus. It will also be pivotal for the identification of potential recombination events between genotypes. Furthermore, it will help identify and analyze viral adaptation events and phenotypic changes associated with specific lineages. It will also guarantee that future studies addressing molecular epidemiology of ZIKV will use standardized and consistent terms across reports and, therefore, allow for an easier sharing of findings among different studies. As such, we consider the adoption of this classification system a crucial step for the future of ZIKV research.

---

**S1 Fig.**
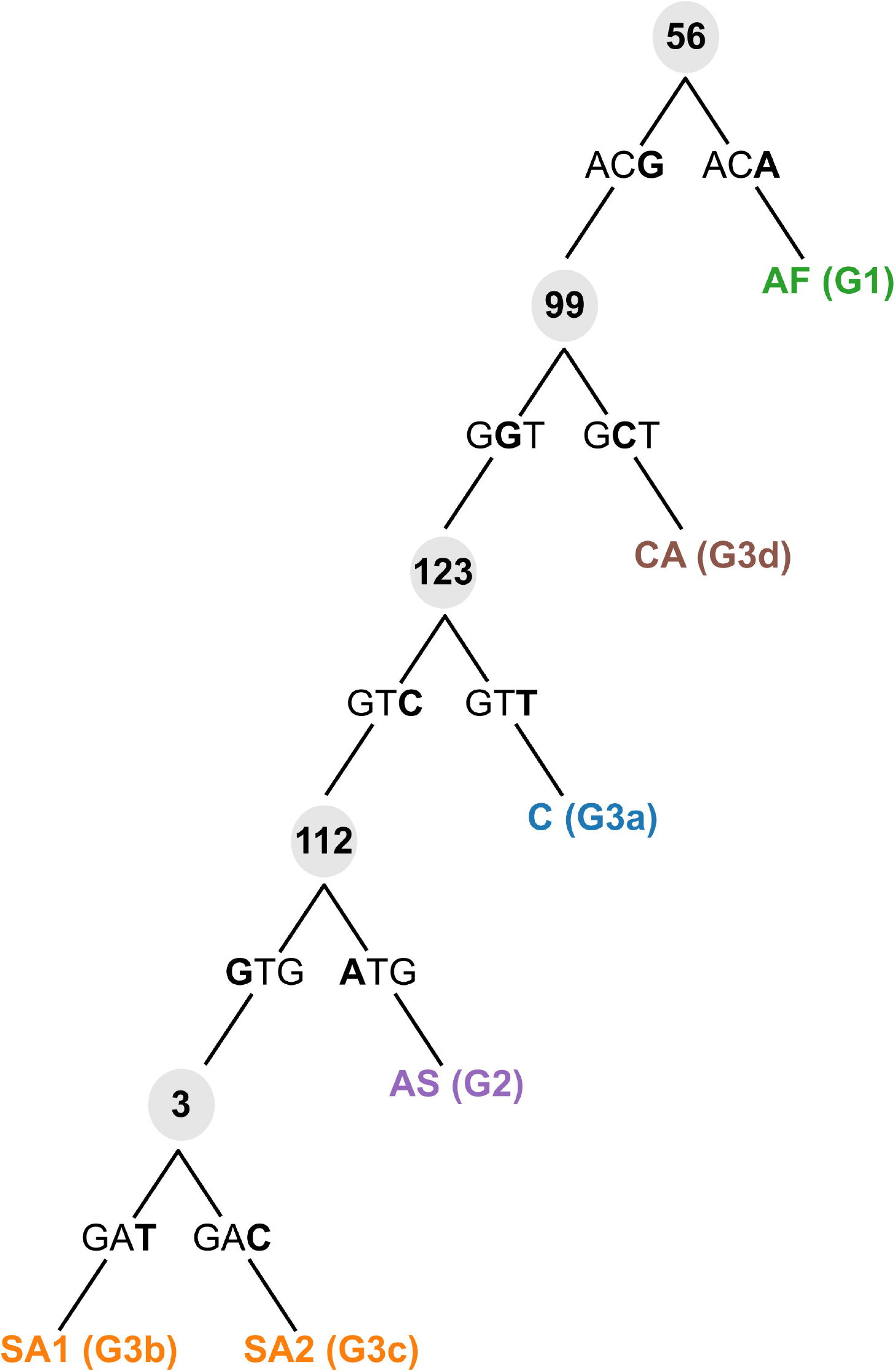
Detailed maximum likelihood trees (IQTree), including bootstrap support, using alignments: 1 (Whole Gnome Sequence), 2 (genome segment 1-4000bp), 3 (genome segment 4001-8000bp), 4 (genes NS1-NS2) and 5 (gene NS3). Color coding denotes geographic origin of strains. Branch labels summarize information about each strain in the format of: “Genbank ID | Country of origin | Collection year”.

**S2 Fig.**
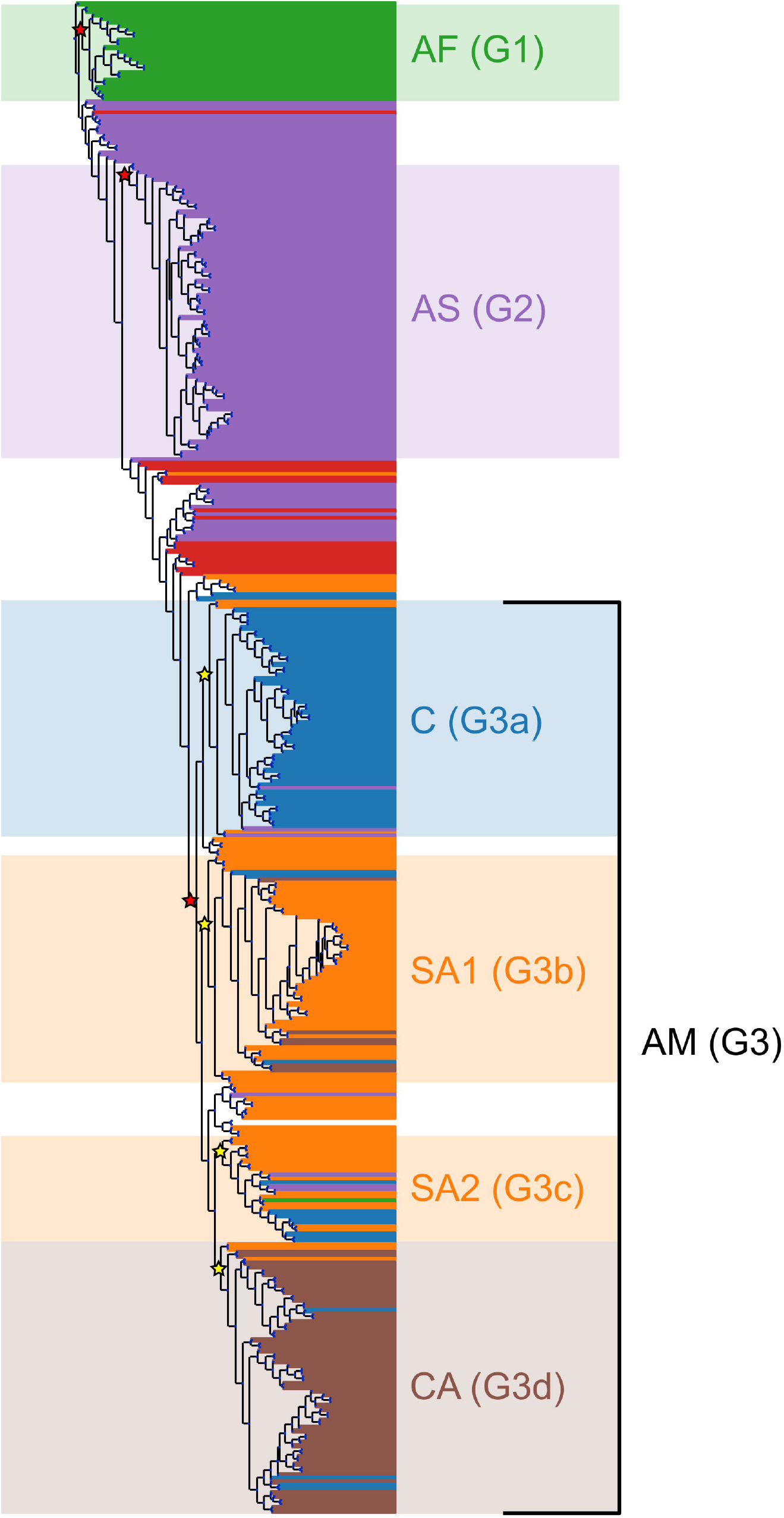
Detailed maximum likelihood tree, including bootstrap support, of whole genome sequences of full dataset (414 strains) using the GTR model in FastTree. Color-coding and branch labelling match that of S1 Fig.

