## Supplementary figures and images for "An updated genotype classification system for Zika viruses"

### SFig1.pdf

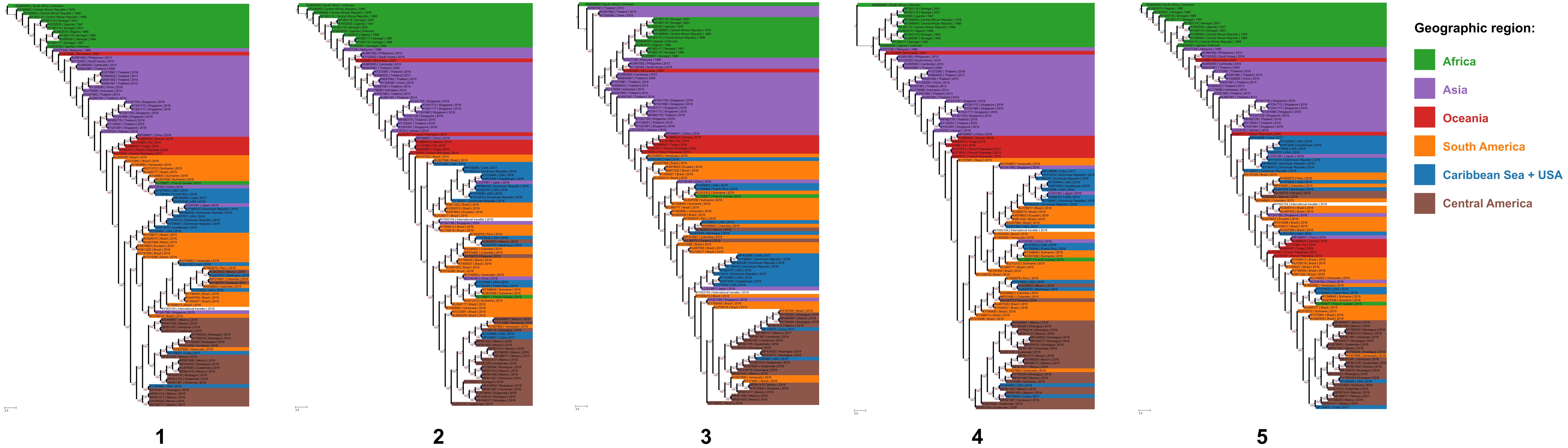

### SFig2.pdf

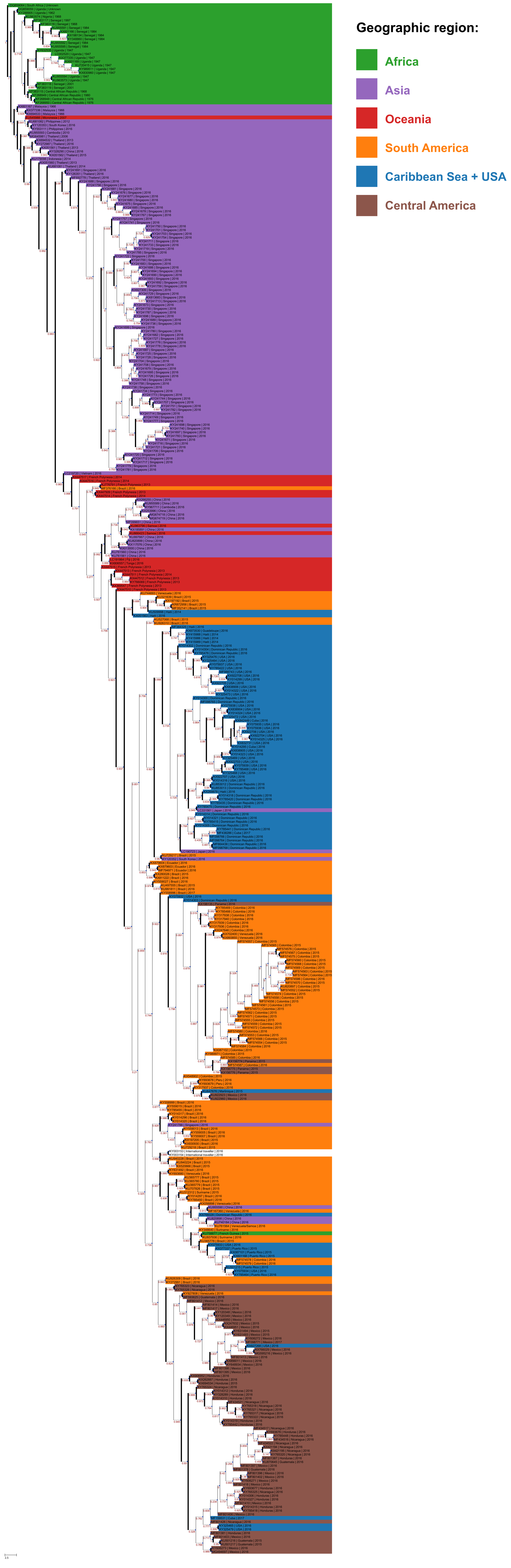
